# Diverse transcriptomic and mutational patterns but limited functional pathway alterations in patient-derived SS cells

**DOI:** 10.1101/2025.03.07.641995

**Authors:** Fernando Gallardo, Evelyn Andrades, Arnau Iglesias, María Maqueda, Teresa Lobo-Jarne, Jessica González, Joan Bertran, David Conde, Eva Rodriguez, Beatriz Bellosillo, Ramon M. Pujol, Anna Bigas, Lluís Espinosa

## Abstract

**Background:** Eradication of SS is hampered by its genetic and molecular heterogeneity. A better understanding of the putative commonalities underlying SS oncogenicity may help to provide more efficient therapeutic strategies against this devastating disease.

**Purpose:** The present work analyzes the whole transcriptome of different patient-derived SS cells to identify expression patterns, functional programs and expressed gene mutations that may provide clues on new therapeutic options for SS patients

**Methods:** Mononuclear cells were recovered by Ficoll gradient separation from fresh peripheral blood of SS patients (n=7). Selected pathway-based compounds and the MALT1 inhibitor MI2 were used for in vitro drug sensitivity testing. SS cells viability was evaluated using CellTiter-Glo_3D Cell Viability Assay and flow cytometry analysis. We validated the usefulness of MI2 using patient-derived SS cells xenotransplanted (PDX) into Nod Scid Gamma mice.

**Results:** In vitro data indicated that cell lines and primary malignant SS cells all display different sensitivities against specific pathway inhibitors. However, MALT1 inhibition led to a robust effect in vitro that was partially reproduced in the in vivo NSG mice xenograft model.

**Conclusions:** Our investigations revealed the actual possibility of inhibiting the downstream TCR signaling complex form by CARD11, BCL10 and MALT1 in SS therapy.

**Key Points:** Patient-derived SS cells are transcriptionally and mutationally heterogeneous but share some common pathway alterations.

Inhibition of MALT1 reduces NF-κB signaling and cell growth in cell lines and patient-derived SS cells.

## INTRODUCTION

Cutaneous T-cell lymphomas (CTCL) comprise a heterogeneous group of non-Hodgkin T-cell malignancies that arise from skin specific T cells and may involve lymph nodes, viscera and peripheral blood. The most frequent CTCL are mycosis fungoides (MF) and the Sézary syndrome (SS)^1^.

While some patients with CTCL have an indolent clinical course, that can be managed with a combination of oral and skin-directed treatments, those with extensive nodal, visceral or blood involvement as well as SS cases are treated with systemic immuno- or biomodulatory-based therapies including immunotherapy-related treatments targeting the surface receptors CD30 (brentuximab) and CCR4 (mogamulizumab) or conventional chemotherapy regimens for mature T-cell neoplasms and allogeneic haematopoietic stem cell transplantation in selected cases^2^.

However, advances in the genetic and molecular knowledge of CTCL tumor cells have not been translated yet into the implementation of targeted therapies with curative response rates, likely associated with the high heterogeneity observed in the genomic analyses, so responses rates to targeted treatments can be achieved in only 30% of patients being often very short-lived associated in part with the high inter- and intra-tumoral heterogeneity of the disease^3^.

Molecular characterization, including deep sequencing studies (Next Generation Sequencing-NGS) of MF and SS, has been performed by several groups in the recent past years, including ours, leading to the identification of multiple tumor-driver mutations, copy number variations or gene fusion which are supposed to contribute to the development of MF/SS by modulating T cell activation, inhibiting apoptosis, activating specific cell function pathways, affecting chromatin remodeling, DNA damage response, cell cycle or immune surveillance impairment^4–10^. Moreover, results from multiple small cohorts have been combined to identify rare mutations and key disease pathways ^11,12^. In general, human CTCL cells have overlapping genomic features, with recurrent alterations that are shared by other mature T-cell neoplasms, including adult T-cell leukemia/lymphoma, angioimmunoblastic T-cell lymphoma, anaplastic large cell lymphoma, nasal-type natural killer/T-cell lymphoma and PTL not otherwise specified^13^.

Thus, it is essential to obtain new personalized therapies and management strategies (precision medicine) adapted to the mechanisms of tumorigenesis present in individual CTCL patient and tumors. This includes the identification of more active therapeutic agents and targeted therapies with mechanisms of action adapted to individual tumor characteristics, well-tolerated and presenting durable responses. The therapeutic responses from clinical trial results with targeted therapies used as monotherapy against altered molecular pathways indicate that sequence or drug combinations schedules that regulate the different damaged sites will be necessary. In this regard, the design and development of preclinical in vitro and in vivo models offers a practical approach.

NF-κB pathway is a well-known tumor promoter in many types of cancers including colorectal, prostate, breast, T acute lymphoblastic leukemia and CTCL among others. In B and T cells, the CARD11/BCL10/MALT1 (CBM) complex initiates NF-κB and p38 signaling in response to B- and T-cell receptor stimulation^14,15^. NF-κB signaling upon receptor activation leads to proliferation and survival and cytokine expression of T cells (reviewed in^16^). Mutations in several NF-κB regulatory elements have been consistently detected in human CTCL including the TNFα receptor 2, TRIM38, CARD11 and MALT1^5,17^, suggesting that NF-κB could represent a therapeutic target for SS patients.

## RESULTS

### SS cells show divergent transcriptional patterns that converge in a subset of functional programs

We previously characterized our in-house cohort of SS patients by the differential response of their malignant T lymphocytes to targeted therapies^18^. By RNA-seq analysis, we investigated the putative heterogeneity at the transcriptomic level of 7 of these patient-derived SS samples (Supplementary Table S1). Principle component analysis (PCA) of RNA-seq data demonstrated the absence of any significant clustering between human SS samples (Figure 1A). Similar heterogeneity was observed in the unsupervised clustering analysis of genes differentially expressed among SS samples (Figure 1B), and we did not observe a significant association between the reported sensitivity of individual SS patient samples^18^ and pathway overrepresentation. However, we detected a general high prevalence of proinflammatory pathways including TNF_NF B^19^, IL6_STAT3^20^ or IL2_STAT5^21^, metabolic regulators such as MYC^22^ or MTORC1^23^ and T cell differentiation pathways such as NOTCH^24^ or HEDGEHOG^25^ (Figure 1C).

**Figure 1.**
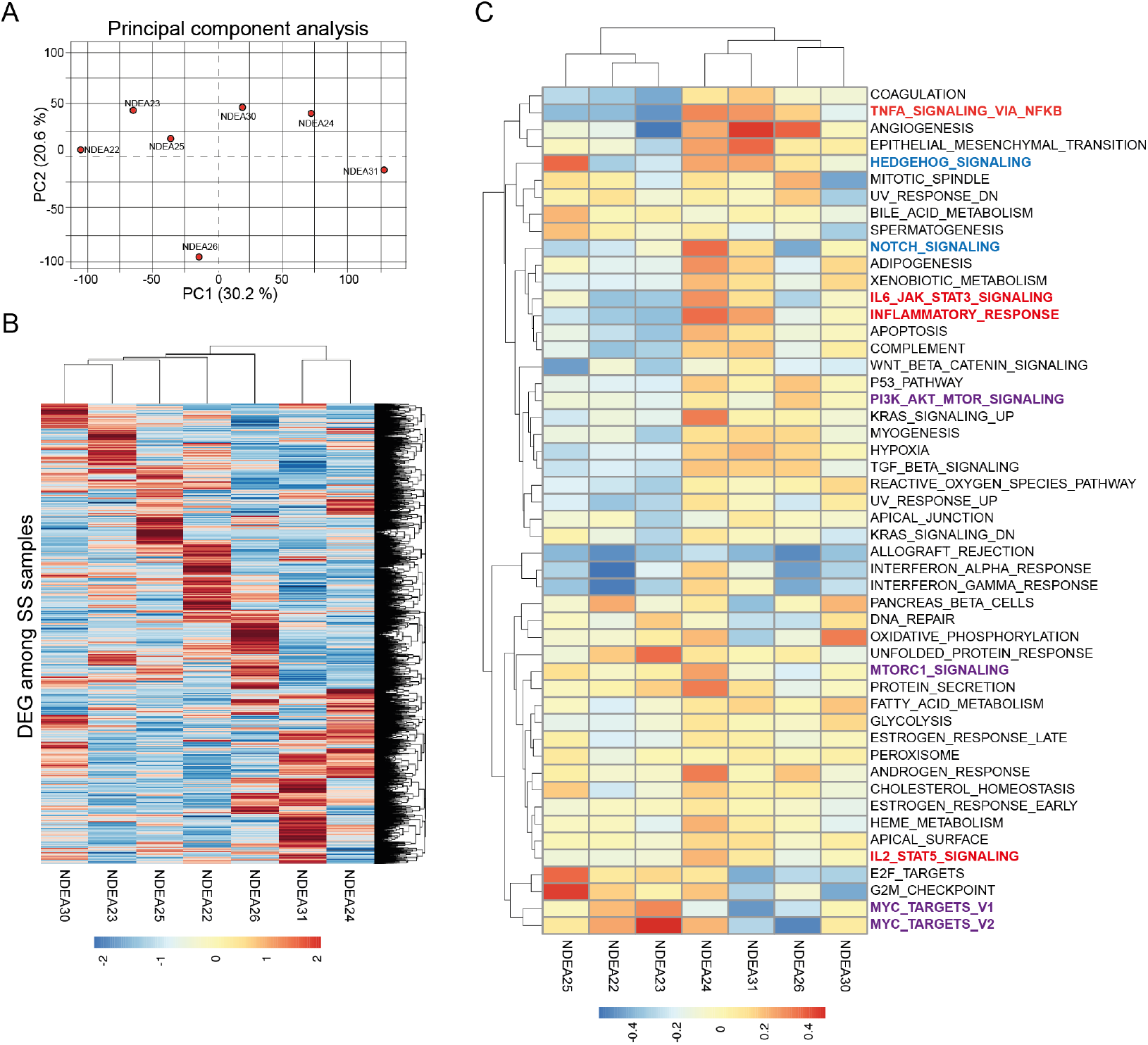
SS samples show divergent transcriptional patterns leading to restricted functional programs. principal component analysis of the RAN-seq data from the 7 SS samples analyzed. Notice the high dispersion observed between samples (A). Heatmap analysis of the same RNA-seq data showing the levels of the differential expressed genes (B) and the pathways corresponding to these genes (C). Notice the convergence of pro-inflammatory (bold red), metabolic regulatory pathways (bold purple) and differentiation pathways (bold blue) in several of the SS samples.

These results indicate that SS cells from individual patients exhibit extensive transcriptional heterogeneity, but share similar features, including the upregulation of common inflammatory, metabolic, and differentiation pathways.

### Mutated genes that are expressed in SS cells include elements of the NF-B pathway and several chromatin-editing enzymes

Several groups have previously investigated the mutational landscape of SS cells by DNA sequencing^5,7,17^. We now examined whether mutated genes are expressed and detectable in the transcriptome of SS cancer cells. After applying stringent filters, including DP (depth of coverage) >=30, allele frequency greater >=0.1 and detection in at least four of the seven samples, we identified an average of 75 mutations per sample, representing a total of 106 mutated genes (Supplementary Table S2 and Figure 2A), which is in concordance with the number of mutated genes per sample (average=51.8) identified by whole exome sequencing in Da Silva Almeida et al^5^. It is noteworthy that only nine of the genes identified in our analysis were common to those identified by Da Silva Almeida (Figure 2B). However, this concordance was even higher than that observed when comparing genomic mutations between samples using comparable filters (Figure 2C). In particular, in our analysis of the Da Silva Almeida data, we detected only two samples (SRR277822 and SRR277812) with four mutations in common (Figure 2C). Among the mutated genes detected by expressed sequence analysis, we identified the NF-B regulatory elements BCL10, CHUK or TRIM38, and DNA/chromatin editing enzymes such as KMT2C and NCOR (Figure 2D). Importantly, TRIM38, NCOR, and KMT2C were among the nine genes identified in common in our and Da Silva Almeida’s analysis (Figures 2B).

**Figure 2.**
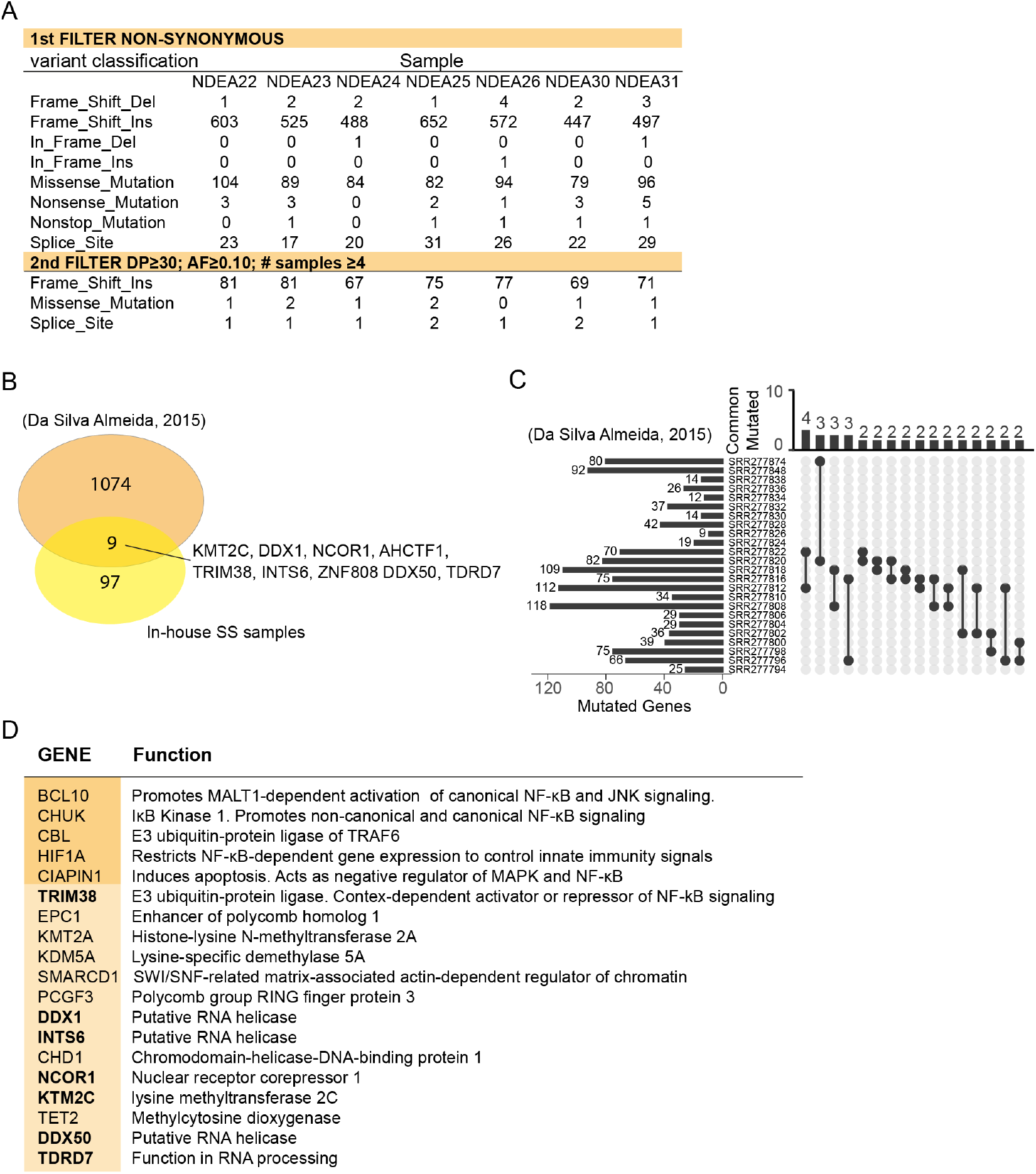
Mutated genes expressed in SS cells are different from sample to samples but they correspond to few functional groups including chromatin editing enzymes and NF-B. Table showing the number and type of mutations identified in the RNA-seq of the SS samples, and the filters used to select actual mutations (A). Venn diagram showing the overlap between genes identified as mutated in our in-house samples and those identified in Da Silva-Almeida et al (B). Analysis of the genes identified as commonly mutated in the different samples analyzed in Da Silva-Almeida et al (C). Examples of genes identified as mutated in several of the samples analyzed corresponding to NF-B regulators, chromatin editing enzymes and RNA helicases (D).

These results indicate that a significant proportion of the genes that are mutated in the SS cells are present in their transcriptome, including several NF-B regulatory elements and chromatin-editing enzymes.

### MALT inhibition reduces NF-B signaling and SS cell growth, and synergizes with multiple targeted therapies in vitro

BCL10 is a core component of the CBM complex, together with the MALT1 paracaspase and CARD11, which drives NF-B and p38/MAPK signaling downstream of the B- and T-cell receptors (reviewed in ^14,15^). MALT1 inhibitors such as MI2 have been proved for their efficacy to eradicate chronic lymphocytic leukemia ^26^ and activated B cell-like diffuse large B cell lymphoma cells ^27^. We aimed to investigate the putative effect of MALT inhibition on NF-B signaling in multiple Cutaneous T-cell Lymphoma (CTCL), which is associated with TAK1 activation^28^. Treatment of CTCL cell lines with the MALT inhibitor MI2 reduced NF-B activation, as indicated by the decrease in the amount of phosphorylated (active) TAK1 and I Bα (Figure 3A). Most importantly, and consistent with the role of NF-κB as a pro-survival and proliferation factor in T-cell leukemias^29,30^, MI2 treatment as a single agent resulted in dose-dependent inhibition of cell growth in all CTCL cell lines tested (Figure 3B) and patient-derived SS samples (Figure 3C). We then tested whether MI2 could potentiate the effect of targeted therapies in CTCL cell lines and SS patient samples. We found a variable response of CTCL cells (Figures 3D, 3E, S3A and S3B) and SS patient samples to the combination treatment with clear synergistic effects in the case of SeAx cells (Figure 3E and 3F) and patient #24 in the double treatment with MI2 and panobinostat. We then tested the effect of MI2 in T-acute lymphoblastic leukemia (TLL) cell lines, both as a single agent and in combination with the standard treatment cocktail including vincristine, dexamethasone and L-asparaginase (VDL). We found that MI2 treatment potentiated the effect of VDL therapy in CEM and JURKAT cells (Figure S3C), which was associated with inhibition of TAK1 phosphorylation/activation (Figure S3D). In contrast, MI2, which did not inhibit TAK1 phosphorylation in RPMI cells, did not potentiate VDL effects in this cell line.

**Figure 3.**
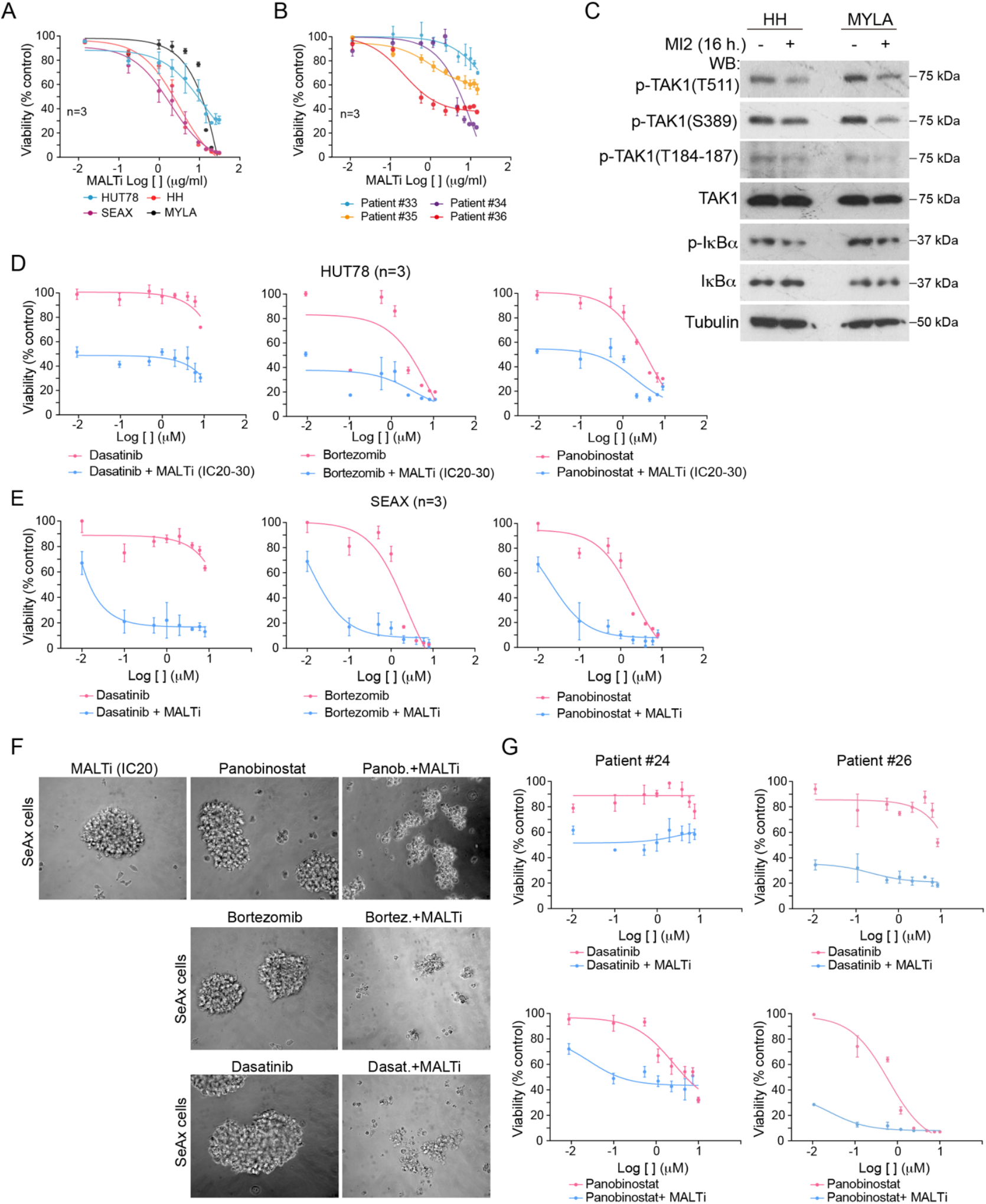
MALTi reduces CTCL cell growth independently of the response to targeted therapies. Dose-response assays of the indicated CTCL cell lines (A) and SS patient samples (B) with the MALT1 inhibitor MI2. Western blot analysis of HH and MYLA cell lines untreated or treated with MI2 for 24 hours to determine the effect of the inhibitor on the NF-B downstream markers p-TAK1 and p-I Bα (C). Dose-response assays of HUT78 (D) and SeAx (E) cell lines with the indicated combination treatment. Microscopic images of SeAx cells treated for 48 hours as indicated (F). Dose-response assays of the primary SS sample from patients #24 and #26 treated as indicated (G).

These results strongly suggest that MALT inhibitors represent a therapeutic option for the treatment of human SS and could be incorporated into combination treatments for selected T-ALL cases.

### MALT inhibition reduces SS cell growth in vivo as single agent

In our in vitro analysis, MI2 was shown to efficiently eliminate most patient-derived SS cells as a single agent, while showing variable effects in combination with targeted therapies (Figure 3B and 3G). We tested the putative therapeutic effect of MI2 on SS cells using a patient-derived xenograft model in NSG mice that we established previously^18^ (Figure 4A). In brief, we transplanted 8 mice with 2×10^6^ SS cells (per mice) from patient #26, which were moderately sensitive to panobinostat but extremely sensitive to MI2 (see Figure 3G). Four weeks after transplantation, we confirmed the presence of SS cells in the peripheral blood (PB) of the recipients by flow cytometry (FC) analysis of human CD45+; CD3+ cells. Since we found slight differences in the number of SS present in each transplanted mouse, the animals were homogeneously assigned to the different experimental groups (vehicle or MI2). After a first round of treatment (intraperitoneal injection for 5 consecutive days and 2 days untreated, repeated twice), animals were left untreated for 7 days before starting a second round of treatment (same protocol) to mimic the procedure used in patients (see treatment procedure in Figure 4B). We performed FC analysis every two weeks to monitor the presence of human CD3+ in the PB of the mice. We detected a significant increase in the number of human CD3+ cells in the PB of both the control and MI2-treated groups after the first round of treatment and the resting period. However, while half of the control mice died during the second treatment and the other mice progressed, none of the MI2-treated animals died during the study period and showed a decrease in the percentage of human CD3+ cells in circulation after the second treatment period (Figure 4C).

**Figure 4.**
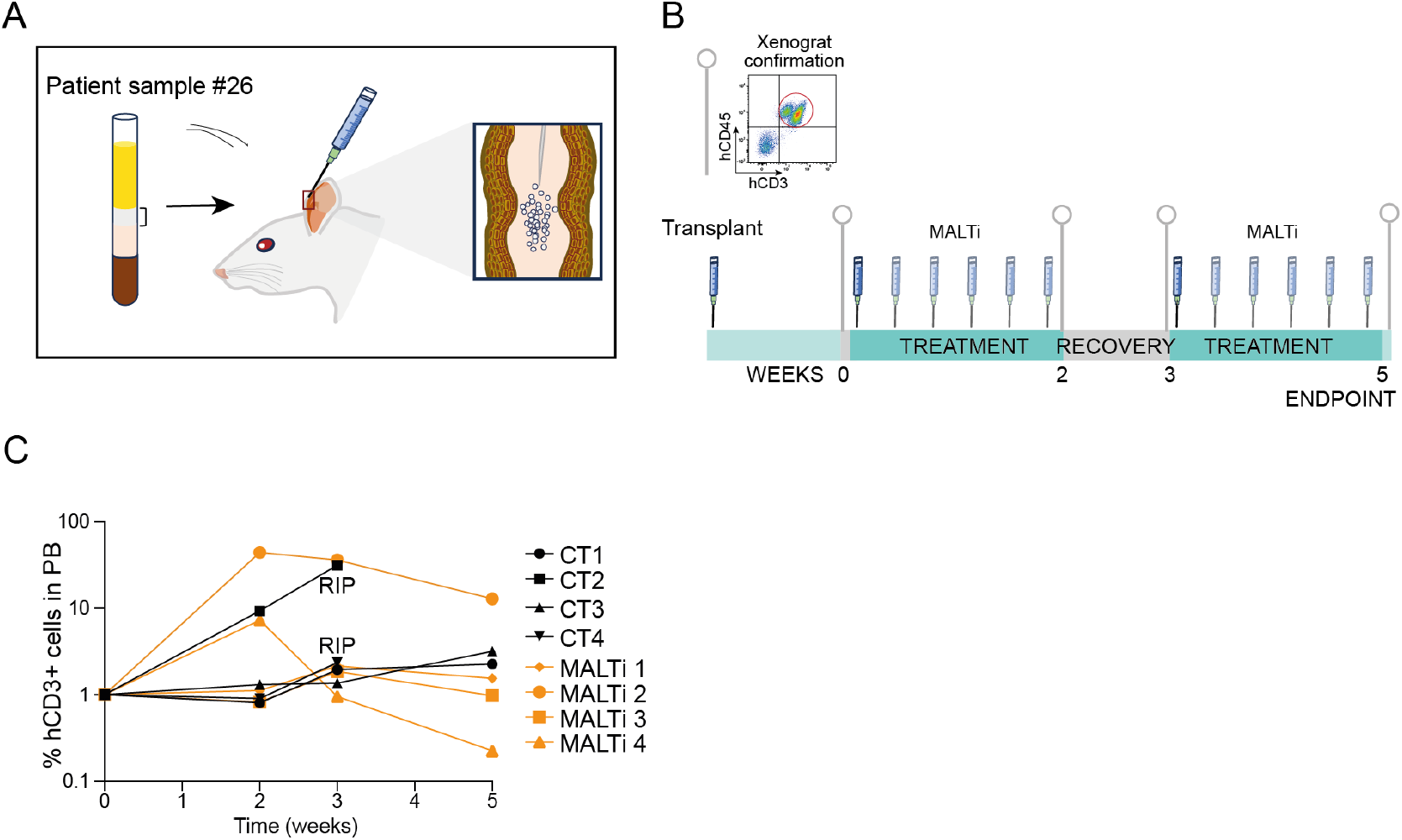
In vivo treatment of SS xenografts. Graphical representation of the procedure used for transplanting the human SS cells obtained from photopheresis (A) and the protocol used in the in vivo testing of treatment efficacy (B). Flow cytometry analysis to determine the proportion of human T cells in the blood stream of the transplanted mice. Notice that 2 of the 4 animals treated with the vehicle died (RIP) during the resting period (C).

Collectively, our data indicate that NF-B signaling acts as a tumor driver in SS cells and can be attenuated by inhibition of the TCR downstream complex formed by CARD11, BCL10 and MALT1 (CBM).

## DISCUSSION

As mentioned, SS is a highly heterogeneous disease both at the level f the cellulat populations involved (highly heterogeneous markers) and at the levels of the mutational background. However, when analyzed considering the pathways to which the mutated elements belong to, there is some association with pathways involved in inflammation, metabolism or organismal development. Particularly, mutated o deregulated elements are found to belong to pathways such as NF-κB or JAK/STAT, and epigenetic regulators^4,11,12,18^. However, the fact that the number of mutations identified in tumor from single patients is very high and are rarely shared between patients have made difficult to propose targeted therapies for SS. These results may indicate the occurrence of a tremendous intratumoral heterogeneity with clones differentially represented in the bulk of the tumor and showing divergent mutational patterns being some of them currently investigated as candidate therapeutic targets in clinical trials for MF or SS, i.e. the PI3K regulator duvelisib or NF-κB inhibitors as well. The HDAC inhibitors vorinostat, romidepsin and resminostat are approved for their use in CTCL by regulatory drug agencies in US and EU. Finally, JAK/STAT inhibitors are increasing its use in different hematological and skin conditions and activators of the antitumor immune response (anti-PD1/PDL1) seem to be positioning as alternative therapeutics in clinical trials ^3,31,32^.

MALT1 inhibitors have been broadly proposed as treatment for B-Cell lymphomas carrying mutations in elements of the CBM complex such as BCL10 or MALT1. Few publications have already suggested that MALT1 could be mutated in SS cells leading to aberrant NF-B activation^33^. However, since of the mutations found consist of nucleotide insertions, whether the alterations are real or artifacts of the detection method has been under debate. We here detected mutations in several elements of the NF-κB pathway including BCL10 in most of the SS samples tested using restricted parameters with an allelic frequency superior or equal to 10%. Most importantly, we experimentally demonstrated that SS cells both cell lines and patient-derived samples are highly sensitive to MALT1 inhibitors showing a variable degree of dependency of NF-κB signaling through CBM. These results are also in agreement with the identification of NF-κB as one of the pathways commonly activated in SS cells. We propose that a combination of MALT1 inhibitors with targeted therapies could represent a realistic treatment for SS patients.

## METHODS

### Patients and study samples

Seven Sézary Syndrome (SS) patients were included in the study. Clinical characteristics, peripheral blood flow cytometry (FC) parameters, patient treatments and clinical status are represented in Supplementary Table S1 and previously described^18^. Primary SS cells used for in vitro drug screening and in vivo generation of PDX were obtained during the apheresis procedure currently done as adjuvant treatment for SS patients. An enriched sample of mononuclear cells was obtained by Ficoll separation. Monitoring and quantification of aberrant malignant SS cells in the patient samples was done by FC for the standardized SS markers CD3+/CD4low/CD26-/CD7-^34–36^ using FITC/PE-conjugated monoclonal antibodies (BD Biosciences, San Diego, CA, USA).

### In vitro drug screening platform and response testing

Small molecules inhibitors used in the study were obtained from Selleck Chemicals drug library (http://www.selleckchem.com/screening/fda-approved-drug-library.html) (Houston, TX, USA). Selection of single-agent testing was based on our previous observations^18^ and included dasatinib (multiple kinase inhibitor), bortezomib (proteasome and NF-B inhibitor), panobinostat (HDAC inhibitor) and MI2 were prepared at 10 mM DMSO stocks and then diluted as indicated. In these experiments, 4×10^5^ cells per well were seeded in 96-well plates and treated with the therapeutic agents for 72 hours. We included additional controls that were treated with the minimum and maximum doses of DMSO used as vehicle (0.1% and 20%). The effect of pharmacological inhibition on cell viability after 72 hours of incubation was determined with CellTiter-Glo_3D Cell Viability Assay (Promega, Madison, WI, USA).

The MALT1 inhibitor MI2 (SS7429) was used in vivo at a concentration of 25mg/Kg according to previous publications^37^.

### In vivo murine model and quantification of therapeutic response

Sézary cells transplantation was performed by intradermal injection of 2 million mononuclear cells in 20uL of PBS (obtained from apheresis) in the ear of NOD-SCID-gamma (NSG) mice (The Jackson Lab, Bar Harbor, ME USA) of 8-12 weeks. PDX assays were carried out in accordance with the regulations of Parc de Salut Mar-Parc de Biomèdica de Barcelona-Universitat Autònoma de Barcelona (TG-09-1223P3-ABS) and the current regulations on Protection of Personal Data and Guarantees of Digital Rights (General Data Protection Regulation of the EU, Regulation EU-2016/679 and Spanish Organic Law 3/2018) and the Biomedical Research Law 14/2007 and Royal Decree RD1716/2011.

First bleeding was performed at 4 weeks after transplantation and subsequent repeated every 2 weeks. We have established detection of 1-10 % of human CD3+ cells in circulation as the criterion to start the in vivo treatment with the MALT1 inhibitor MI2. After starting treatment, mice were bled weekly to quantify by FC the human T-cell lymphoid populations. At sacrifice, either at the end-point of the study or when mice should be euthanized because of disease progression, we analyzed tumor burden in PB by FC.

## ACKNOWLEDGEMENTS

We would like to thank all members of Drs. Gallardo, Espinosa and Bigas labs from their support and helpful comments and suggestions. This work was supported by Instituto de Salud Carlos III and co-funded by the European Union (PI22/00069 and PI21/0390) and grant 2017-SGR 135 from AGAUR from the Government of Catalonia and CIBERONC.

## AUTHORSHIP CONTRIBUTIONS

FG, AB and LE designed the study, analyzed data and wrote the manuscript.

EA, AI, MM, TL-J, JG, BB and JB performed experiments and analyzed data.

DC, ER and RM-P provided reagents, analyzed data and revised the manuscript.

## Supplementary Figures and Legends

**Figure S1.**
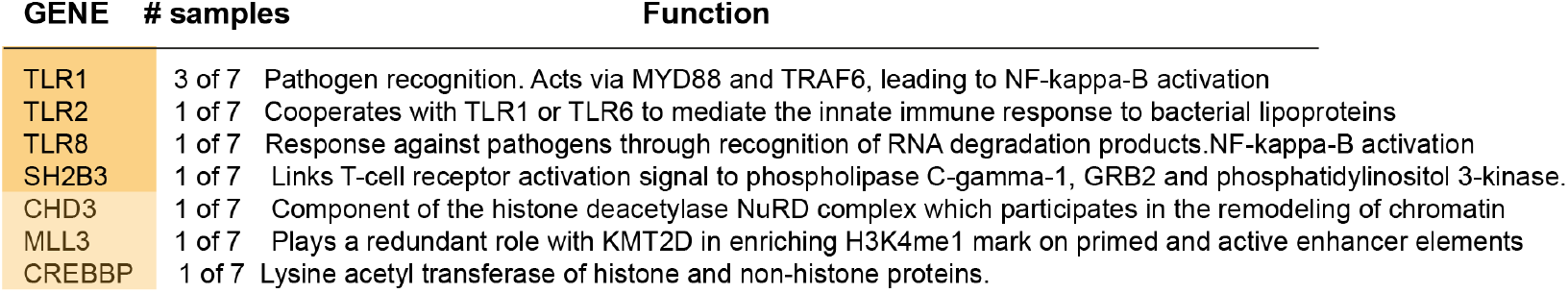
Other mutated genes expressed in SS cell correspond to few functional groups including chromatin editing enzymes and NF-B signaling.

**Figure S2.**
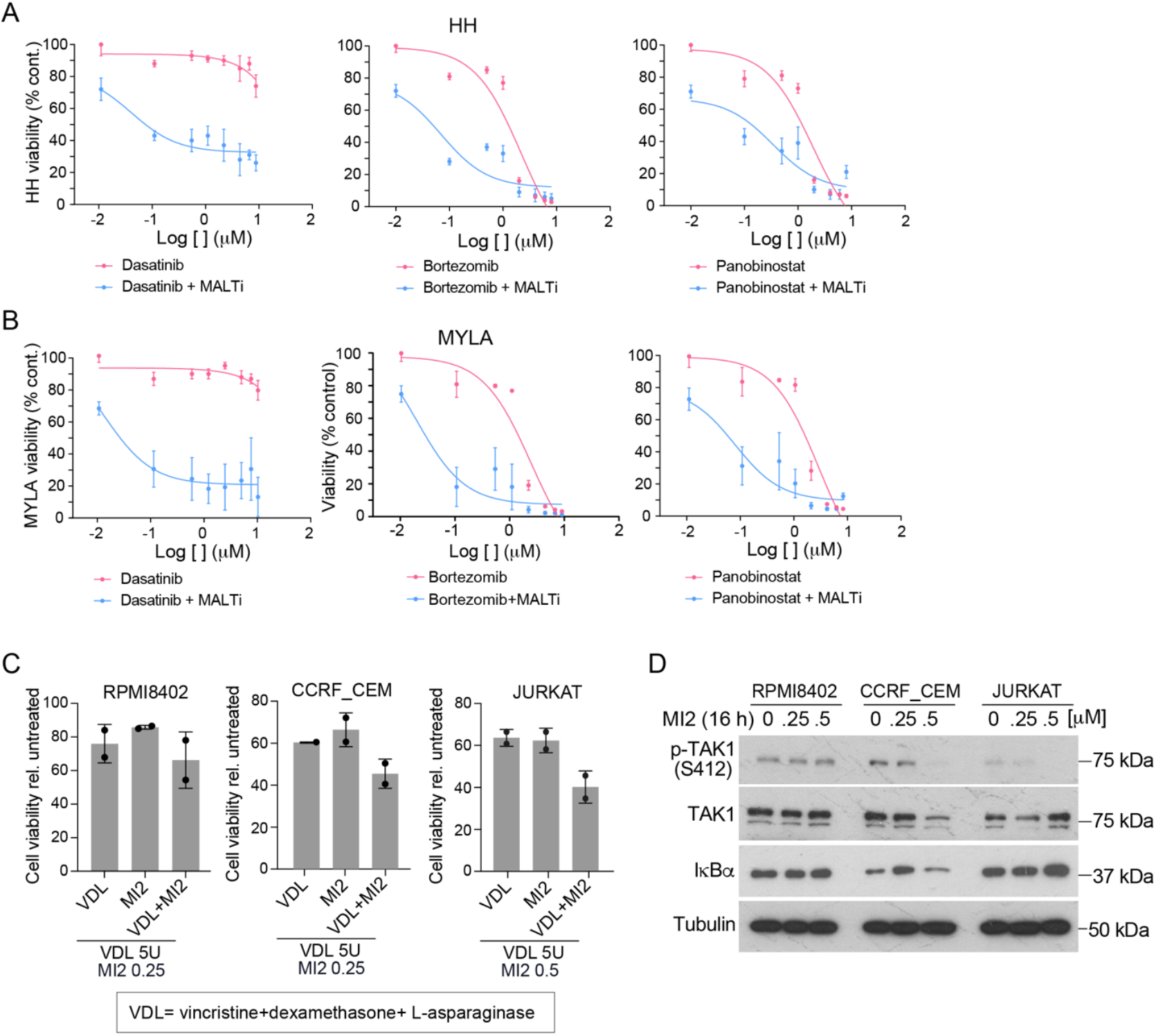
Synergistic effect of MALTi with multiple targeted therapies. Dose-response assays of HH (A) and MYLA (B) CTCL cells treated as indicated. Effects of MI2 treatment on the viability of the indicated T-ALL cell lines, either as monotherapy or in combination with the standard VDL treatment (C). Western blot analysis to show the effects of MI2 treatment in TAK1 activation (phosphorylation of serine 412).

## References

1. Willemze, R. et al. The 2018 update of the WHO-EORTC classification for primary cutaneous lymphomas. Blood 133, 1703–1714 (2019).

2. Latzka, J. et al. EORTC consensus recommendations for the treatment of mycosis fungoides/Sézary syndrome - Update 2023. Eur. J. Cancer 195, 113343 (2023).

3. Quaglino, P. et al. Phenotypical Markers, Molecular Mutations, and Immune Microenvironment as Targets for New Treatments in Patients with Mycosis Fungoides and/or Sézary Syndrome. J. Invest. Dermatol. 141, 484–495 (2021).

4. Tensen, C. P., Quint, K. D. & Vermeer, M. H. Genetic and epigenetic insights into cutaneous T-cell lymphoma. Blood 139, 15–33 (2022).

5. da Silva Almeida, A.C. et al. The mutational landscape of cutaneous T cell lymphoma and Sézary syndrome. Nat. Genet. 47, 1465–1470 (2015).

6. Prasad, A. et al. Identification of Gene Mutations and Fusion Genes in Patients with Sézary Syndrome. J. Invest. Dermatol. 136, 1490–1499 (2016).

7. Wang, L. et al. Genomic profiling of Sézary syndrome identifies alterations of key T cell signaling and differentiation genes. Nat. Genet. 47, 1426–1434 (2015).

8. Woollard, W. J. et al. Candidate driver genes involved in genome maintenance and DNA repair in Sézary syndrome. Blood 127, 3387–3397 (2016).

9. Vaqué, J. P. et al. PLCG1 mutations in cutaneous T-cell lymphomas. Blood 123, 2034–43 (2014).

10. Caprini, E. et al. Loss of the candidate tumor suppressor ZEB1 (TCF8, ZFHX1A) in Sézary syndrome. Cell Death Dis. 9, 1178 (2018).

11. Park, J. et al. Genomic analysis of 220 CTCLs identifies a novel recurrent gain-of-function alteration in RLTPR (p.Q575E). Blood 130, 1430–1440 (2017).

12. Chang, L.-W. et al. An Integrated Data Resource for Genomic Analysis of Cutaneous T-Cell Lymphoma. J. Invest. Dermatol. 138, 2681–2683 (2018).

13. Palomero, T. et al. Recurrent mutations in epigenetic regulators, RHOA and FYN kinase in peripheral T cell lymphomas. Nat. Genet. 46, 166–70 (2014).

14. Lucas, P. C., McAllister-Lucas, L. M. & Nunez, G. NF-kappaB signaling in lymphocytes: a new cast of characters. J. Cell Sci. 117, 31–9 (2004).

15. Thome, M. & Tschopp, J. TCR-induced NF-kappaB activation: a crucial role for Carma1, Bcl10 and MALT1. Trends Immunol. 24, 419–24 (2003).

16. Hayden, M. S. & Ghosh, S. NF-κB in immunobiology. Cell Res. 21, 223–44 (2011).

17. Ungewickell, A. et al. Genomic analysis of mycosis fungoides and Sézary syndrome identifies recurrent alterations in TNFR2. Nat. Genet. 47, 1056–1060 (2015).

18. Gallardo, F. et al. Sézary syndrome patient-derived models allow drug selection for personalized therapy. Blood Adv. 6, 3410–3421 (2022).

19. Karin, M. NF-B as a Critical Link Between Inflammation and Cancer. Cold Spring Harb. Perspect. Biol. 1, a000141–a000141 (2009).

20. Tanaka, T., Narazaki, M. & Kishimoto, T. IL-6 in inflammation, immunity, and disease. Cold Spring Harb. Perspect. Biol. 6, a016295 (2014).

21. Surbek, M., Tse, W., Moriggl, R. & Han, X. A centric view of JAK/STAT5 in intestinal homeostasis, infection, and inflammation. Cytokine 139, 155392 (2021).

22. Schaub, F. X. et al. Myc-Directed Suppression of Autophagy Provides Therapeutic Vulnerabilities Targeting Amino Acid Homeostasis. Blood 126, 2450–2450 (2015).

23. Panwar, V. et al. Multifaceted role of mTOR (mammalian target of rapamycin) signaling pathway in human health and disease. Signal Transduct. Target. Ther. 8, 375 (2023).

24. Brandstadter, J. D. & Maillard, I. Notch signalling in T cell homeostasis and differentiation. Open Biol. 9, 190187 (2019).

25. Crompton, T., Outram, S. V & Hager-Theodorides, A. L. Sonic hedgehog signalling in T-cell development and activation. Nat. Rev. Immunol. 7, 726–35 (2007).

26. Saba, N. S. et al. MALT1 Inhibition Is Efficacious in Both Naïve and Ibrutinib-Resistant Chronic Lymphocytic Leukemia. Cancer Res. 77, 7038–7048 (2017).

27. Fontan, L. et al. MALT1 small molecule inhibitors specifically suppress ABC-DLBCL in vitro and in vivo. Cancer Cell 22, 812–24 (2012).

28. Gallardo, F. et al. Novel phosphorylated TAK1 species with functional impact on NF-κB and β-catenin signaling in human Cutaneous T-cell lymphoma. Leukemia 32, 2211–2223 (2018).

29. Vilimas, T. et al. Targeting the NF-kappaB signaling pathway in Notch1-induced T-cell leukemia. Nat. Med. 13, 70–7 (2007).

30. Espinosa, L. et al. The Notch/Hes1 Pathway Sustains NF-κB Activation through CYLD Repression in T Cell Leukemia. Cancer Cell 18, 268–281 (2010).

31. Zinzani, P. L. et al. Panoptic clinical review of the current and future treatment of relapsed/refractory T-cell lymphomas: Peripheral T-cell lymphomas. Crit. Rev. Oncol. Hematol. 99, 214–227 (2016).

32. Trautinger, F. et al. European Organisation for Research and Treatment of Cancer consensus recommendations for the treatment of mycosis fungoides/Sézary syndrome – Update 2017. Eur. J. Cancer 77, 57–74 (2017).

33. Willis, T. G. et al. Bcl10 is involved in t(1;14)(p22;q32) of MALT B cell lymphoma and mutated in multiple tumor types. Cell 96, 35–45 (1999).

34. Feng, B. et al. Flow cytometric detection of peripheral blood involvement by mycosis fungoides and Sézary syndrome using T-cell receptor Vβ chain antibodies and its application in blood staging. Mod. Pathol. 23, 284–295 (2010).

35. Horna, P. et al. Flow cytometric evaluation of peripheral blood for suspected Sézary syndrome or mycosis fungoides: International guidelines for assay characteristics. Cytom. Part B Clin. Cytom. 100, 142–155 (2021).

36. Pulitzer, M. P., Horna, P. & Almeida, J. Sézary syndrome and mycosis fungoides: An overview, including the role of immunophenotyping. Cytom. Part B Clin. Cytom. 100, 132–138 (2021).

37. Goral, A. et al. A Specific CD44lo CD25lo Subpopulation of Regulatory T Cells Inhibits Anti-Leukemic Immune Response and Promotes the Progression in a Mouse Model of Chronic Lymphocytic Leukemia. Front. Immunol. 13, 781364 (2022).

