## Supplementary Table 1 for "Diverse transcriptomic and mutational patterns but limited functional pathway alterations in patient-derived SS cells"

| Patient | Age | Gender | Stage | Sézary Cell Count; % Leukocytes | CD3+/CD4+/CD7-/CD6- (% of T Lymphocytes) | Status | More sensitive drugs |
|---|---|---|---|---|---|---|---|
| #23 | 64 | M | IVA1 | $15.18x10^9$/L; 36 | 88%* | Alive-PR | Bortezomib, Bardoxolone Methyl |
| #25 | 63 | M | IVA1 | $1.10x10^9$ /L; 5.4 | 45% | Alive-SD | Bortezomib, Sorafenib, Temsirolimus, Bardoxolone Methyl, Dasatinib, Vorinostat, Doxorubicin, Gemcitabine, Neralabine, Aprepitant |
| #22 | 65 | M | IVA1 | $1.13x10^9$/L; 22 | 86% | Alive-WOD | Bortezomib, Sorafenib, Bardoxolone Methyl, Aprepitant |
| #24 | 71 | F | IVA2 | $1.05x10^9$/L; 30 | 95% | Alive-SD | Bortezomib, Bardoxolone Methyl, Panobinostat |
| #26 | 64 | M | IVA1 | $1.52x10^9$/L; 29 | 89% | Alive-PR | Bortezomib, Temsirolimus, Bardoxolone Methyl, Doxorubicin, Aprepitant |
| #30 | 77 | F | IVA1 | $4.17x10^9$/L; 25 | 69% | Alive-SD | Bortezomib |
| #31 | 81 | F | IVA1 | $3.04x10^9$/L; 24 | 71% | Alive-SD | Bortezomib |
| #32 | 66 | M | IVA1 | $14.77x10^9$/L; 59 | 86% | Alive-SD | Belinostat, Panobinostat, Romidepsin |

SD: stable disease
PR: partial remission
WOD: without disease
*(associated abnormal 19% circulating B-cell population)
